# ProAgio, a Novel Integrin αvβ3 Targeted Cytotoxin, Suppresses Tumor Growth and Reprograms the PDAC Microenvironment

**DOI:** 10.64898/2026.01.15.699725

**Authors:** Dhana Sekhar Reddy Bandi, Sujith Sarvesh, Ganji Purnachandra Nagaraju, Harrison Kim, Jeremy Foote, Sejong Bae, Karina J. Yoon, Midhun Malla, Ashiq Masood, Mehmet Akce, Zhi-Ren Liu, Bassel F. El-Rayes

**Author notes:** **Corresponding Author: Bassel F. El-Rayes, MD**, Albert F. LoBuglio Endowed Chair for Translational Cancer Research, Division Director, Hematology and Oncology, Deputy Director, O’Neal Comprehensive Cancer Center, Heersink School of Medicine, University of Alabama at Birmingham.

## Abstract

**Background:** Pancreatic ductal adenocarcinoma (PDAC) is characterized by a dense, hypoxic and immune-suppressive tumor microenvironment (TME). Integrin αvβ3-expressing cells, including endothelial and cancer-associated fibroblasts (CAFs), contribute to the development of this TME. ProAgio is a novel cytotoxin that targets integrin αvβ3-expressing cells. ProAgio is currently in clinical trials. We have previously shown that the combination of GPH (gemcitabine, paricalcitol, and hydroxychloroquine) influences PDAC TME. Based on the overlapping mechanisms of action, we hypothesized that ProAgio could potentiate effects of GPH and enhance its anti-tumor immunity.

**Methods:** Patient-derived xenograft (PDX) and orthotopic models of PDAC were used to assess the therapeutic activity and mechanism of action of ProAgio in combination with GPH. Immunohistochemistry was used to evaluate hypoxia, EMT and angiogenesis. Changes in the immune cells were measured with multi-parameter flow cytometry. Dynamic contrast-enhanced MRI (DCE-MRI) was used to study tumor perfusion in mice and patients (NCT06182072).

**Findings:** ProAgio potentiated the growth inhibitory effects of GPH in PDX and orthotopic models by depleting integrin β3 expressing cells, leading to ECM remodeling, reduced vascular leakage, improved hypoxia, and reversed EMT. DCE-MRI showed a significant increase in tumor perfusion following ProAgio treatment in mice and patients (NCT06182072). Immune profiling revealed that the combination treatment significantly increased the infiltration of γδ T cells, natural killer T (NKT) cells, CD4^+^ effector T cells, and M1-like macrophages. Furthermore, the combination treatment reduced the expression of myofibroblastic CAFs (myCAFs), further supporting the immunomodulatory and stromal normalizing effects of GPH and ProAgio.

**Conclusion:** Targeting integrin αvβ3 using ProAgio modulates the PDAC TME by improving perfusion, reducing hypoxia, reversing EMT, and alleviating immune suppression. ProAgio potentiates the effects of GPH therapy, which should be evaluated in future trials.

## Introduction

Pancreatic ductal carcinoma (PDAC) resists immune and cytotoxic therapies ^1^. The tumor microenvironment (TME) of PDAC is characterized by dense desmoplastic stroma, aberrant angiogenesis, hypoxia, high prevalence of immune suppressive cells, and lack of anti-tumor effector immune cells, all of which contribute to the resistance to therapy. Cancer-associated fibroblasts (CAFs) deposit ECM, a central PDAC TME feature. CAF secrete immunosuppressive cytokines, limiting the recruitment and activation of anti-tumor effector immune cells. ^2^ In response to the hypoxic TME, PDAC cells synthesize and secrete proangiogenic cytokines such as vascular endothelial growth factor (VEGF) ^3^. The dense desmoplasia and persistent elevation in proangiogenic factors lead to the development of immature endothelial cells and leaky blood vessels that, in turn, contribute to increased intestinal pressure, decreased blood flow, and hypoxia ^4, 5^. These changes in the TME impair drug delivery ^6^. The hypoxic TME also promotes the development of resistance to therapy through epithelial to mesenchymal (EMT) changes, autophagy activation, and immunosuppressive environment developmentent. Simultaneous targeting of activated CAF and aberrant angiogenesis in PDAC can modulate TME, resulting in decreased hypoxia, increased perfusion and drug delivery, ultimately reversing resistance to therapy ^7–9^.

Integrin α_v_β_3_is expressed in activated CAFs and immature endothelial cells. Our group developed a protein-based drug, ProAgio, an integrin α_v_β_3_-targeted cytotoxin. ProAgio binds to a non-catalytic site of β_3_ protein (ITGB3), activating caspase pathways and apoptosis ^10^. In preclinical models, ProAgio depleted integrin α_v_β_3_ expressing CAFs and endothelial cells, leading to enhanced activity of cytotoxic chemotherapy. ProAgio also reduced the levels of insulin growth factor-1 by depleting CAF stroma, which in turn reduced cytidine deaminase and increased gemcitabine-induced apoptosis ^11, 12^. ProAgio is currently in clinical trials in combination with gemcitabine and nab-paclitaxel (NCT06182072) in metastatic PDAC.

Our group previously published a comprehensive study demonstrating enhanced antitumor effects of gemcitabine (G) and nab-paclitaxel by hydroxychloroquine (H) and paricalcitol (P) ^13^. The combination induced autophagy. In addition, GPH favorably modulated the TME by promoting the transition of activated CAFs to quiescence, increasing the intratumoral M1 to M2 ratio, CD4^+^, CD8^+^, and NK cells. Furthermore, in a clinical trial, these effects were observed in paired biopsies from patients with PDAC treated with P, H, G, and nab-paclitaxel (<u>NCT04524702</u>). Based on the mechanism of action of PH and ProAgio and the nonoverlapping toxicities, we proposed investigating the combination of GPH with ProAgio in PDAC. In this study, we hypothesize that targeting integrin α_v_β_3_-expressing CAF and endothelial cells, combined with the effects of PH on CAFs and PDAC cells, will reduce hypoxic niches in the tumor microenvironment, enhance perfusion, lower resistance, and improve efficacy of immune and cytotoxic chemotherapy.

### Dynamic contrast-enhanced (DCE) MRI

DCE-MRI was used to evaluate vascularity changes in PDAC following ProAgio treatment in both the orthotopic KPC mouse model and human subjects. In the animal study, group 1 (n=3) received intraperitoneal injections of ProAgio (15 mg/kg) daily for six days, while group 2 (n=3) received saline injections. Small animal MRI imaging was conducted using a Bruker BioSpec 9.4T system (Bruker BioSpin Corp., Billerica, MA), 0, 2, and 6 days after therapy initiation. A T1 map was obtained using a multi-flip-angle gradient-echo sequence with the following settings: TR/TE = 115/3 ms, 128×128 matrix, 30×30 mm field of view, NEX=4, and flip angles of 10°, 20°, 30°, 40°, 50°, 60°, and 70°. DCE-MRI was performed using the same parameters as the T1 mapping sequence, with a fixed flip angle of 30°. Gadoteridol (0.0267 mmol/ml) was injected via a syringe pump (NE-1600, New Era Pump Systems, Inc., Wantagh, NY) at a constant rate of 0.01 ml/sec, delivering a total volume of 0.15 ml. A lab-made computer software based on LabVIEW (National Instruments Co., Austin, TX) was employed to calculate the volume transfer constant (*K^trans^*) using the reference region mode ^14^. Comparisons of *K^trans^*changes were performed using one-way repeated measures ANOVA.

Our group is currently conducting a phase Ib dose escalation clinical trial (NCT06182072) evaluating the safety of ProAgio with gemcitabine and nab-paclitaxel in patients with stage IV PDAC. Gemcitabine (1000 mg/m^2^ I.V.) and nab-paclitaxel (100mg/m^2^ I.V.) were administered on days 1, 8, and 15. ProAgio is administered at 6.4 mg/Kg I.V. weekly in the first dose level and 10.7 mg/kg I.V. weekly in the second dose level. Based on prior pharmacokinetic studies, the ProAgio dose of 6.4 mg/Kg is sub-therapeutic, while the 10.7 mg/kg provides a therapeutic dose. Patients had research DCE-MRI scans before (baseline) and 8±1 weeks after the start of treatment. Tumor boundary was determined on the CT images by board-certified radiologists blinded to therapy, obtained before and 8±1 weeks after the ProAgio treatment. All participants were instructed to abstain from consuming solid food and caffeinated beverages for at least 24 hrs before imaging. Imaging was performed using a single 3T MRI scanner (GE Signa). All patients were scanned with P4 phantoms for scanner-dependent error correction ^15^. A 3D fast spoiled gradient echo sequence (FSPGR) was employed for DCE-MRI with the following parameters: matrix size = 256 x 230, FOV = 400×360 mm, frequency/phase encoding = 192/173, TR/TE = 3.8/2.1 ms, slice number = 12, thickness/gap = 5/0 mm, SENSE factor = 2, NEX = 1, flip angle = 15°, and temporal resolution = 2.9 seconds. The same imaging parameters were used for T1W mapping, but with three flip angles (2°, 5°, and 10°). T1W imaging for both T1 mapping and DCE-MRI was conducted in a free-breathing mode, and the images obtained at the expiration phase were automatically selected and co-registered using the B-spline method ^16^. Gadoteridol (0.1 mmol/kg) was intravenously administered at a rate of 2 mL/s, starting 30 seconds after the initiation of DCE-MRI, followed by a 20 mL saline flush at the same rate (2 mL/s). The *K^trans^* maps were created using an extended Tofts model with a population-based arterial input function and lab-made computer software based on LabVIEW (National Instruments Co., Austin, TX).

## Results

### ITGB3 is highly expressed in PDAC and correlates with poor prognosis

To evaluate the role of ITGB3 in PDAC, we analyzed its expression using IHC in a human PDAC tissue array (PA483-L97, Biomax) that contains 40 cancer and eight non-cancer tissues (**Supplementary Table 1**). The expression of ITGB3 was significantly higher in PDAC tissues than in non-cancerous pancreas tissues (**Figure 1A-C**). We then used TCGA data to investigate whether the previously observed high expression of ITGB3 correlates with genes involved in hypoxia, EMT, glycolysis, and angiogenesis. To this end, we analyzed 176 patients’ data with a median ITGB3 expression of 3.1 pTPM (protein-coding transcripts per million). Patients were divided into high and low expression groups with 23.5 pTPM and 0.1 pTPM, respectively. High ITGB3 expression was significantly associated with HIF-1α (hypoxia) (**Figure 1D**). Similarly, ITGB3 was highly correlated with GLUT1, linked to metabolic reprogramming (**Figure 1E**). ITGB3 was also positively correlated with angiogenic and vascular remodeling markers such as VEGF-A (**Figure 1F**) and CD31 (**Figure 1G**). High ITGB3 mRNA expression was positively correlated with vimentin (**Figure 1H**). ITGB3 also positively correlated with ECM markers such as COL1A1, a key component in ECM remodeling and tumor-stroma interactions (**Figure 1I**). These findings suggested that ITGB3 is overexpressed at the mRNA and protein levels in PDAC and is associated with EMT, angiogenesis, ECM remodeling, glycolysis, hypoxia and metastasis.

**Figure 1:**
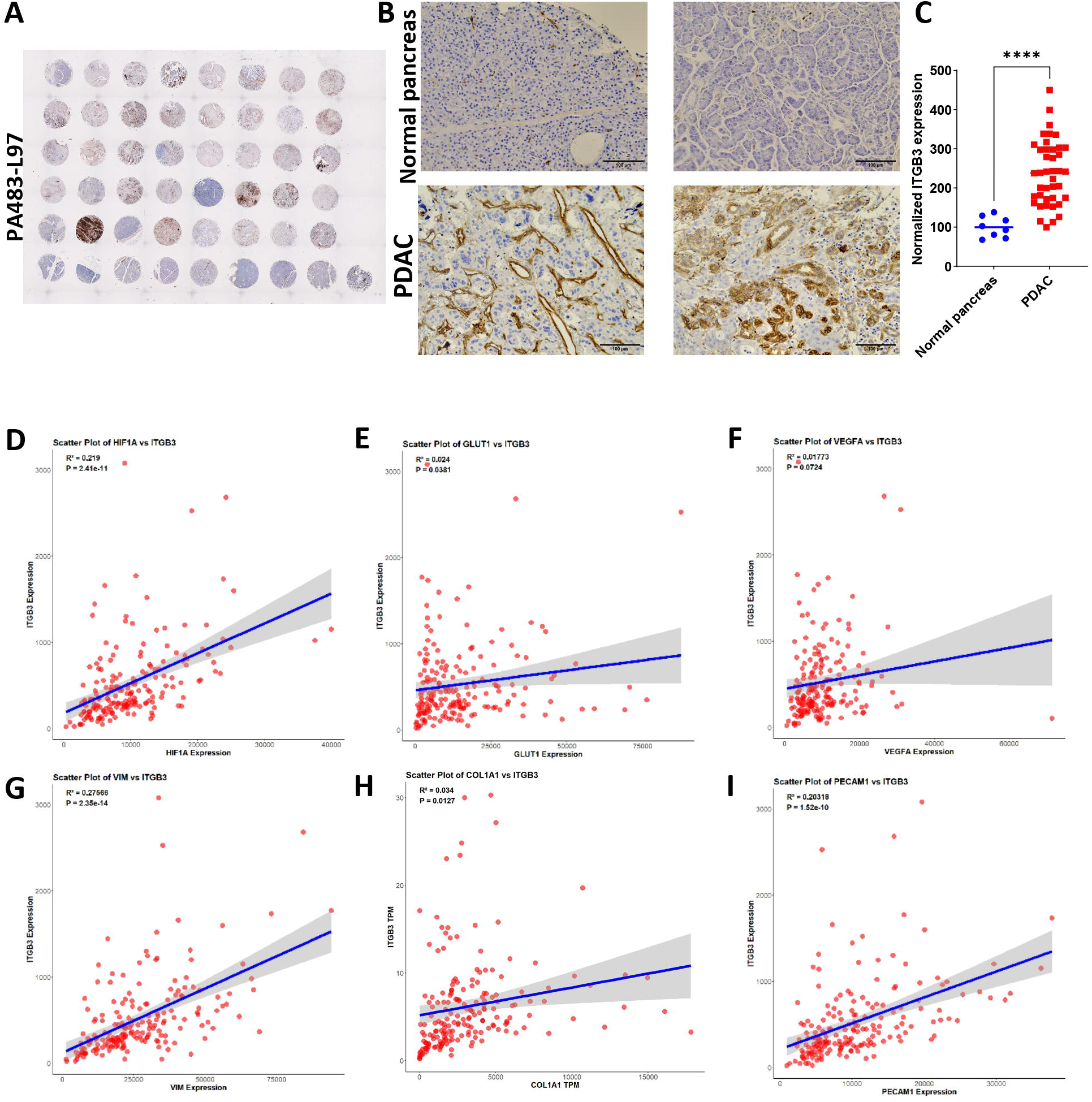
ITGB3 overexpression in human PDAC tissues correlates with poor survival. **(A)** Representative IHC staining of ITGB3 in PDAC tissue microarrays (TMA) showing varying expression levels in non-cancer and cancer tissue sections. **(B).** High magnification IHC images demonstrating the expression of ITGB3 on normal pancreas and cancer cells with stromal regions. **(D-I)**. Scatter plots depicting the correlation between ITGB3 and key tumor-associated markers such as HIF-1α **(D)**, GLUT-1 **(E)**, VEGFA **(F)**, Vimentin **(G)**, COL1A1 **(H)**, and PECAM1 **(I)**. Pearson correlation coefficients (R) and *p*-values indicate statistically significant positive correlation between ITGB3 and these markers. Statistical significance was assessed using a two-tailed unpaired Student’s t-test. All quantitative data represent the mean ± SEM. ∗∗∗∗p < 0.0001.

### The combination of ProAgio and GPH inhibits the growth of PDAC in murine models

Luciferase-labelled KPC (KPC-*f*-luc) with point mutations in KRAS^G12D^ and p53^R172H^ were surgically implanted into the pancreas of C57BL6/J mice. Mice were imaged one week after implantation (day 0) and randomized into four groups (n=5 animals/group) treated with vehicle, ProAgio, GPH, or GPH plus ProAgio as described in the materials and methods section. Treatment effects on tumor growth were measured by IVIS bioluminescence imaging. Mice were euthanized on day 21, and tumor weights were measured. The GPH and ProAgio combination treatment showed significant growth inhibition compared to ProAgio, GPH, or vehicle (**Figure 2A-B**). The wet tumor weights significantly reduced size and weight in GPH and ProAgio treatment (**Figure 2D-E**). No indications of toxicity, such as changes in body weight or organ abnormalities, were observed (**Figure 2C**).

**Figure 2:**
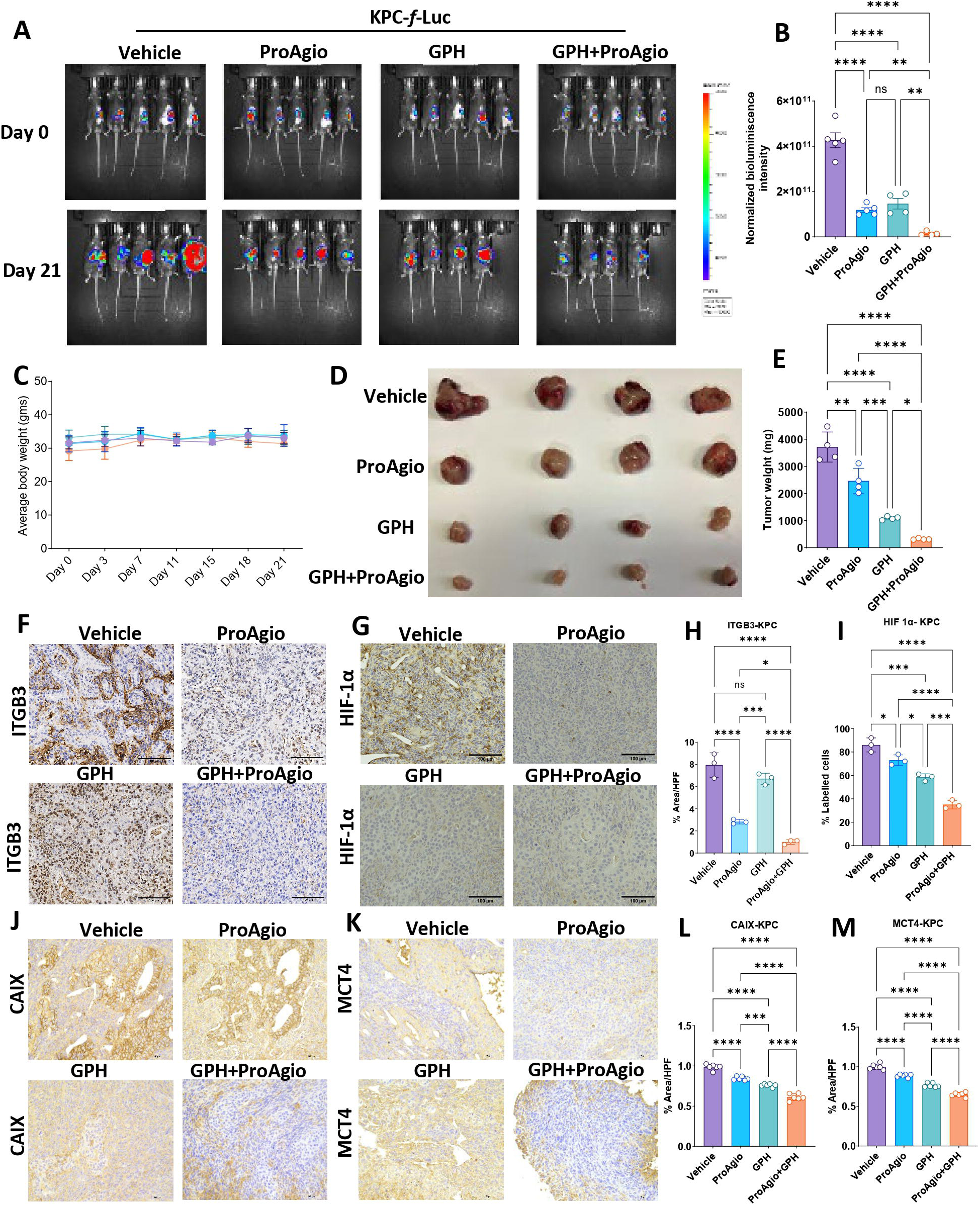
GPH plus ProAgio combination significantly suppresses tumor growth in murine orthotopic PDAC models and reduces hypoxia. **(A).** KPC-*f*-luc-labelled cells were orthotopically implanted into C57BL6/J/J mice (n=20). After 1 week, five mice were orally administered vehicle, ProAgio, GPH, or GPH plus ProAgio. Note: one mouse from the GPH+ProAgio group died before the end of the experiment (n=4 for this group). Bioluminescence images at the indicated time points are shown. **(B).** Normalized bioluminescence values from the panel (A). **(C).** Average body weights of the animals during the treatment are shown in experiment (A). **(D).** Tumor images were taken at the end of the experiment for A. **(E).** Percentage of tumors weights at the indicated conditions for the experiment in (A). One-way ANOVA was used to determine p-values. Error bars indicate SD. ns, non-significant, ∗p < 0.05, ∗∗∗p < 0.001 and ∗∗∗∗p < 0.0001.

### ProAgio plus GPH combination suppresses aberrant angiogenesis, reduces “leaky” blood vessels, and increases perfusion

Integrin α_v_β_3_is expressed on immature endothelial cells involved in aberrant angiogenesis. Using tissue from the KPC orthotopic experiments, IHC analysis revealed significant downregulation in CD31 in GPH plus ProAgio compared to the other treatment groups, suggesting a significant anti-angiogenic effect (**Figure 4A** and **Figure 4J**). We then evaluated if this anti-angiogenic effect was associated with a physiologic change in vascular leakage. We used FITC-Dextran to assess the integrity of blood vessels and evaluate leakage. The combination of GPH plus ProAgio significantly reduced the fluorescent intensity, indicating a reduction in blood vessel permeability compared to vehicle, ProAgio, or GPH (**Figure 3A**). Treatment with GPH or the vehicle exhibited comparatively higher fluorescence levels than ProAgio, further confirming ProAgio’s effects on reducing leaky blood vessels.

**Figure 3:**
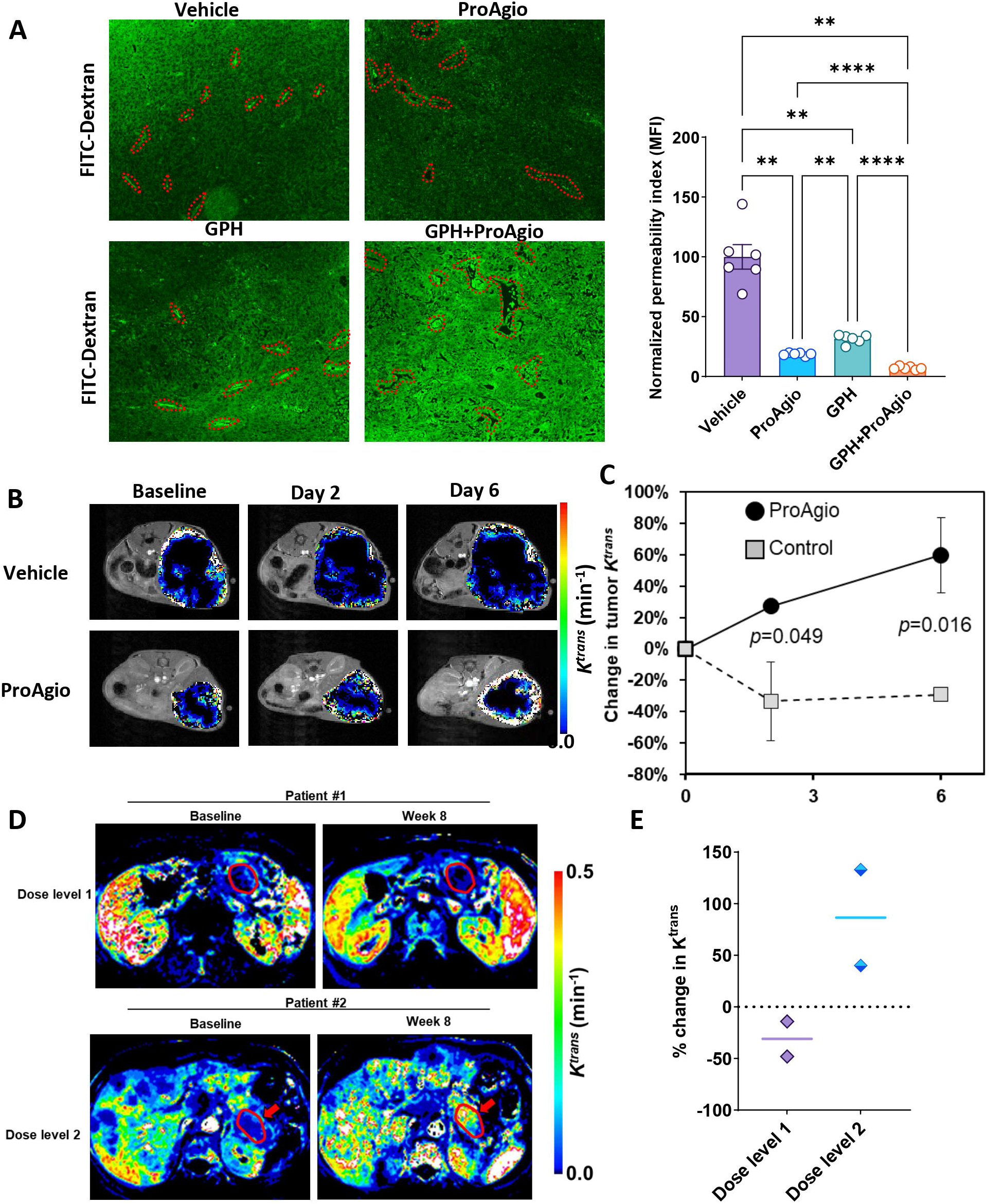
GPH plus ProAgio reduces vascular leakage and enhances drug delivery. **(A).** Functional (or perfusion) changes assessed by the FITC-dextran infusion (70 kDa) show reduced leakage (or permeation of FITC) in GPH plus ProAgio combination compared to single agents or vehicle (left) and the quantifications of permeability index (right). **(B).** K*^trans^*maps of KPC mice treated with untreated (first row) or ProAgio (second row) for over a week. **(C).** The change (%) of averaged K*^trans^* in tumors treated (n=3) or untreated mice (n=3) for a week. The *p*-value is inserted to represent the statistical difference on each measurement day. **(D).** K*^trans^* maps of two subjects having metastatic PDAC showing reduced (1st row) or increased blood flow (2nd row) after 8 weeks of therapy with ProAgio and gemcitabine nab paclitaxel. The tumor region is indicated with a red arrow in each sub-figure. **(E).** Quantifying percentage change in K*^trans^*intensity between baseline and after 8 weeks of treatment with ProAgio gemcitabine and nab paclitaxel. ProAgio was administered at 6.4 mg/Kg I.V. weekly (Dose level 1) and 10.7 mg/week (dose level 2). Each dot represents a unique patient. Statistical significance was assessed using One-way ANOVA. All quantitative data represent the mean ± SEM. ∗∗*p* < 0.01 and ∗∗∗∗*p* < 0.0001.

**Figure 4:**
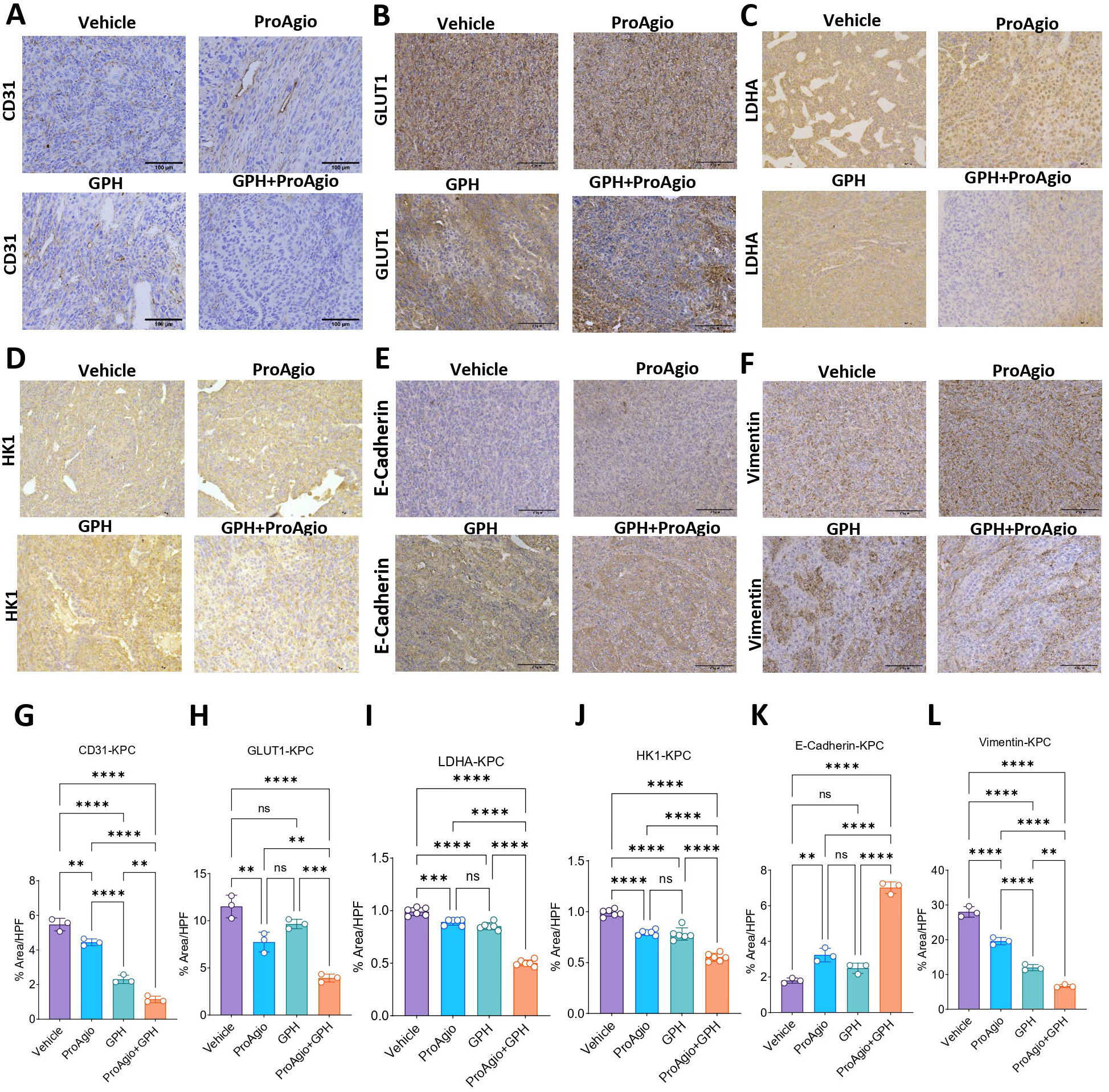
GPH plus ProAgio suppresses angiogenesis, glycolysis, and EMT in PDAC tumors. **(A-I).** Immunohistochemistry (n=5 mice) results showing the reduced expression of CD31 **(A)**, GLUT1 **(B)**, LDHA **(C)**, HK1 **(D)**, E-Cadherin **(E)**, vimentin **(F)**, ITGB3 **(G)**, HIF-1α **(H)**, and CAIX **(I)** in orthotopic KPC-*f*-luc tumors. **(J-R).** IHC quantifications for the the images shown in panel **A-I.** Scale bar 100 µm. Representative IHC quantification is shown in the bottom. One-way ANOVA was used to determine p-values. Error bars indicate SD. ns, non-significant, ∗p < 0.05, ∗∗*p* < 0.01, ∗∗∗p < 0.001 and ∗∗∗∗p < 0.0001.

We performed DCE MRI studies to assess the impact of the anti-angiogenesis and vascular leakage on perfusion. **Figure 3B** presents T2-weighted images overlapped with tumor *K^trans^* maps (color scale) of two representative KPC mice, one group (n=3) treated with ProAgio (bottom row) and the other (n=3) untreated (top row). As summarized in **Figure 3C**, the percentage changes in *K^trans^* values within the tumor region after 2 days and 6 days of treatment were -33±25% and -29±5%, respectively, in the ProAgio-treated group. In contrast, the corresponding changes in the untreated group were 27±2% and 60±24%, respectively. These differences were statistically significant, with *p* values of 0.049 and 0.016, respectively.

In our ongoing Phase Ib trial (NCT06182072), three patients with metastatic treatment-naïve pancreatic adenocarcinoma have been treated on each of the first two dose levels. Paired DCE-MRIs were performed on two patients at each dose level. Two subjects treated on dose level 1 were Caucasian females at the ages of 61 and 62, respectively, and two subjects treated on dose level 2 were a Caucasian female at age 61 and a Caucasian male at the age of 64. **Figure 3D** shows the *K^trans^* maps of the subject treated with a higher dose and a lower dose of ProAgio for 8 weeks. The tumor region is indicated with a red arrow in each sub-figure. The *K^trans^* value of two subjects treated with lower sub-therapeutic doses decreased by 14% and 48%, respectively, for 8 weeks of ProAgio, gemcitabine and nab paclitaxel therapy, whereas that of the subject treated with the higher therapeutic dose increased by 40% and 133% during the same time (**Figure 3D** and **3E)**.

### ProAgio plus GPH combination improves hypoxia, suppresses glycolysis and EMT

ProAgio is an integrin α_v_β_3_-targeted cytotoxin. We evaluated the effects of treatment on ITGB3 expression. ProAgio treatment significantly lowered ITGB3 expression. The combination of GPH plus ProAgio resulted in a significant decrease in ITGB3 compared to monotherapies or vehicles (**Figure 4B** and **Figure 4K**). Given that the ProAgio GPH combination improves perfusion, we investigated its impact on hypoxia. IHC analyzed tumors from the KPC orthotopic mouse experiment for key hypoxia markers, including HIF-1α and CAIX. The combination treatment significantly reduced the expression of HIF-1α (**Figure 4C** and **Figure 4L)** compared to vehicle, ProAgio, or GPH, indicating that the combination therapy. Carbonic anhydrase IX (CA9) is among the highest expressed proteins in response to hypoxia and is a transcriptional target for HIF-1α. GPH plus ProAgio significantly reduced the expression of CA9, suggesting that this combination effectively disrupts hypoxia-driven signaling (**Figure 4D** and **Figure 4M)**. To investigate the impact of combination treatment on glycolytic flux, we first analyzed the expression of GLUT1. We found that combination treatment significantly reduces the expression of this marker, suggesting a potential decrease in glycolytic activity (**Figure 4E** and **Figure 4N**). Next, we analyzed the expression of additional glycolysis-associated markers, including hexokinase 1 (HK1) and lactate dehydrogenase A (LDHA). HK1 catalyzes the first step of glycolysis, and LDHA facilitates the conversion of pyruvate to lactate under anaerobic conditions. IHC analysis revealed that GPH plus ProAgio significantly reduced the expression of HK1 (**Figure 4F** and **Figure 4O**) and LDHA (**Figure 4G** and **Figure 4P**) compared to vehicle tumors, GPH or ProAgio. These changes paralleled the decrease in hypoxia markers, indicating effective blockade of hypoxia-driven glycolysis. The observed reduction in glycolytic markers further strengthens a marked metabolic reprogramming that could make tumor cells more vulnerable to nutrient deprivation and apoptosis. Hypoxic stress is a known driver of EMT and immune exhaustion ^17,18^. The GPH ProAgio combination treatment led to a substantial increase in the expression of E-cadherin (**Figure 4H** and **Figure 4Q**), reduction in vimentin (**Figure 4I** and **Figure 4R**), indicating EMT inhibition and a transition toward a more epithelial and less invasive phenotype.

### GPH plus ProAgio combination alters the immune landscape in the murine PDAC model

We performed multi-parameter flow cytometry to investigate the effects of GPH plus ProAgio on the immune landscape. First, we observed a significant elevation of CD45^+^ immune cells in the combination treatment compared to vehicle, ProAgio, or GPH (**Figure 5A**). This represents a broad population of immune cells, indicating enhanced recruitment and infiltration of these cells into the TME. We then characterized further the subtypes of these immune cells. A concurrent increase in the number of γδ and T NK cells but not NK cells was observed in GPH plus ProAgio (**Figure 5B-D**). Furthermore, the combination therapy increased the frequencies of T NK cells. It increased the expression of the immune checkpoint markers such as PD-1, CTLA4 and TIGIT NK cells, suggesting an activated phenotype (**Figure 5D**).

**Figure 5:**
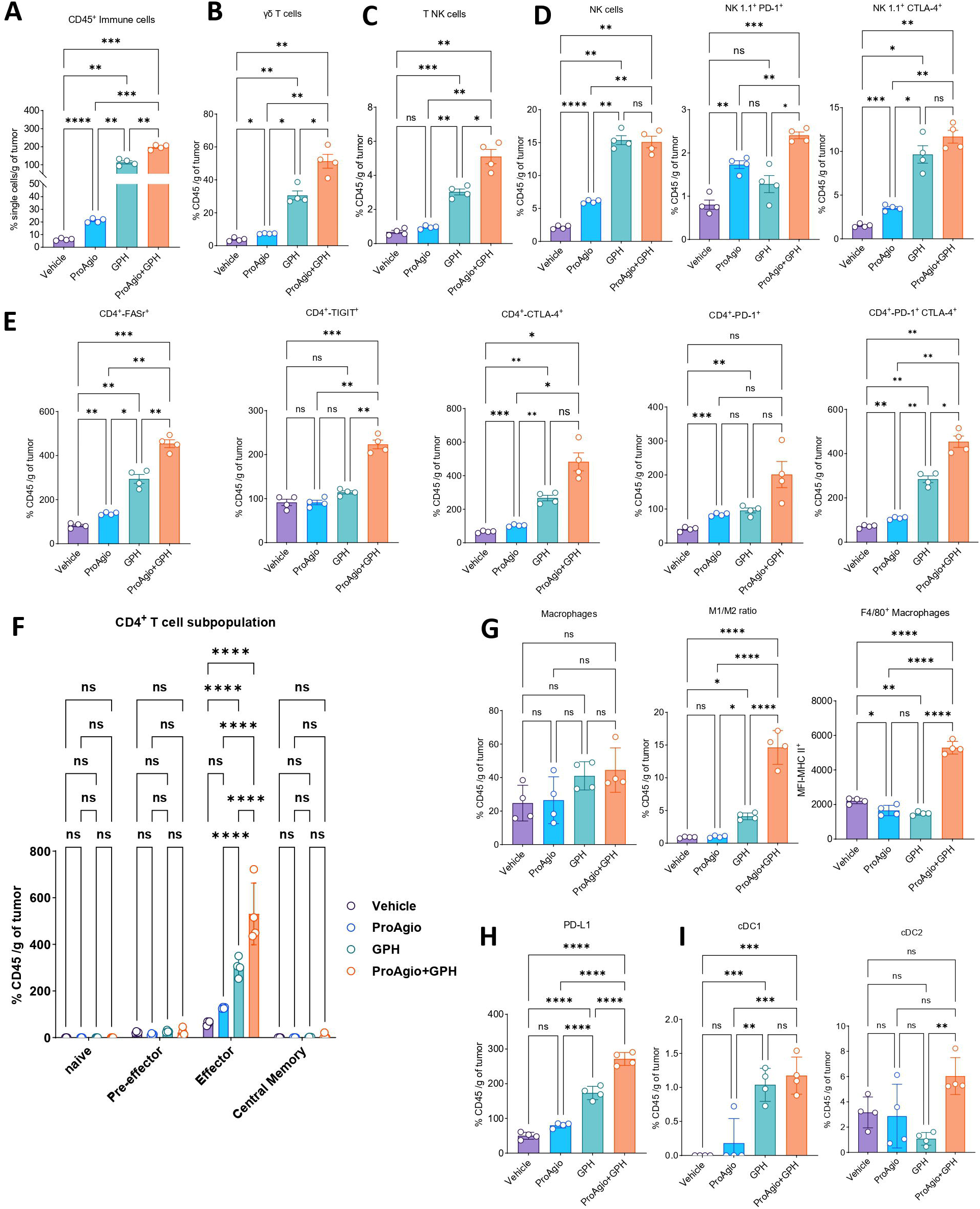
GPH plus ProAgio combination enhances anti-tumor immunity. **(A).** Proportions of CD45^+^ immune cells, **(B).** γδ T-cells, **(C).** TNK cells **(D).** NK cells, NK1.1^+^ PD-1^+^, NK1.1^+^ CTLA-4^+^ cells. **(D).** Quantities of CD4^+^ FASr^+^, CD4^+^ TIGIT^+^, CD4^+^ CTLA-4^+^, CD4^+^ PD-1^+^, and CD4^+^ PD-1^+^ CTLA-4^+^ T cells in KPC TiME in vehicle, ProAgio, GPH, and GPH+ProAgio (n=4) treated mice. (**F**). Expressions of CD4^+^ subset populations such as naïve, pre-effector, effector, and central memory T cells in vehicle, ProAgio, GPH, and GPH+ProAgio. (**G**). expression of macrophages, increased M1/M2 ratio, and the increased percentages of CD45^+^ CD11b^+^ F4/80 in the conditions shown. (**H**). Compared to single agents, increased CD11b^+^ F4/80^+^ PD-L1^+^ percentages were observed in combination treatment. **(I).** Increased cDC1 and cDC2 expressions in GPH+ProAgio compared to vehicle, ProAgio, or GPH-treated mice. One-way ANOVA was used to determine *p*-values. Error bars indicate SD. ns, non-significant, ∗*p* < 0.05, ∗∗*p* < 0.01, ∗∗∗*p* < 0.001, and ∗∗∗∗*p* < 0.0001.

Examining the CD4^+^ T cell population (**Figure 5E**) showed a significant increase in FAS receptor (FASr), TIGIT, CTLA-4, PD-1 and the co-expression of PD-1 and CTLA-4 on CD4^+^ T cells in the combination therapy (**Figure 5E**), suggestive of active priming and expansion of activated CD4^+^ T cells in the combination-treated groups. GPH plus ProAgio monotherapy induced lower levels of these markers with minimal increase compared to the vehicle-treated group. In addition to the increase in immune checkpoint markers on CD4^+^ T cells, we also observed significant changes in the phenotype of CD4^+^ T cell subsets following treatment with GPH plus ProAgio (**Figure 5F**). Notably, there was a significant increase in the effector CD4^+^ T cells, without changes in the frequencies of naïve, pre-effector, and central memory CD4^+^ T cells, suggesting that the combination treatment enhances priming, activation and differentiation of the CD4^+^ T cells (**Figure 5F**). The findings of increased immune checkpoint markers on CD4^+^ T and NK cells suggest that a checkpoint inhibitor may be needed to maximize the immune modulatory effects of GPH plus ProAgio.

Interestingly, while a substantial increase in the immune checkpoint inhibitors was also observed in NK cells, the CD8^+^ T cells did not show a similar trend. There was no significant increase in CD8^+^ T cells (**Supplementary figure 1A**), and CD8^+^ CTLA-4^+^ (**Supplementary figure 1B**), CD8^+^ PD-1^+^ (**Supplementary figure 1C**), CD8^+^ CTLA-4^+^ PD-1^+^ (**Supplementary figure 1D**). In addition to this, the CD8^+^ T cell compartment showed no substantial changes in naïve, pre-effector, effector, and central memory T cells (**Supplementary Figure 1E**). Together, these results highlight the preferential activation of CD4^+^ T cells in response to combination treatment. Possible reasons for this phenomenon would be that alterations in the TME that favor MHC-II antigen presentation, increased APC engagement, or therapy-induced cytokine signaling that promotes helper T cell proliferation could all cause the preferential activation of CD4 T cells ^19–22^.

The combination therapy also increased the expression of MHC-II on F4/80^+^ macrophages, indicating tumor-associated macrophage (TAM) polarization into a pro-inflammatory phenotype to support ongoing adaptive immune compartment priming within the tumor immune microenvironment (TiME) (**Figure 5G**). In addition to macrophages exhibiting enhanced pro-inflammatory phenotype, F4/80^+^ macrophages also showed increased expression of PD-L1 in combination therapy (**Figure 5H**), warranting immune checkpoint therapy to maximize immune cell compartment potential in controlling the tumor growth. Our flow cytometry results showed increased F4/80^+^ macrophages and an elevated M1/M2 ratio in the GPH plus ProAgio combination. To confirm, we examined macrophage infiltration using IHC for CD68. Per the previous results, we see a significant increase in CD68 expression in the combination group compared to vehicle, ProAgio or GPH alone. These results further strengthen the idea that combination treatment promotes macrophage recruitment and polarization toward an M1-like phenotype to a more immunostimulatory TME (**Supplementary figures 3A** and **3C**). In line with the above-mentioned observation, the frequency of the conventional dendritic cell population in combination therapy was significantly higher compared to the respective monotherapy, further bolstering the evidence to support enhanced T cell priming (**Figure 5I**).

### GPH plus ProAgio combination inhibits CAFs

Our previously published research demonstrated that GPH therapy reprograms CAFs substantially to achieve a quiescent phenotype ^13^. Since both GPH and ProAgio affect CAFs, we next evaluated the effects of the combination on CAFs ^11,13^. Multi-parameter flow cytometric evaluation of CAFs revealed a trend towards an increase in myCAFs (CD45^-^EPCAM^-^PDPN^+^LY6C^-^) with vehicle, GPH or ProAgio treatment. GPH plus ProAgio combination reduced myCAFs significantly compared to GPH (**Figure 6A**). As the myCAFs are known to enhance fibrosis, we evaluated ECM remodeling by analyzing COL1A1 expression through IHC. IHC analysis revealed a notable decrease in COL1A1 levels in tumors subjected to combination therapy compared to vehicle, ProAgio, or GPH monotherapies (**Supplementary figure 3B** and **3D)**. Treatment with GPH increased iCAFs (CD45^-^EPCAM^-^PDPN^+^LY6C^+^), whereas ProAgio to GPH reduced iCAFs significantly (**Figure 6B**). GPH plus ProAgio significantly increased apCAFs (CD45^-^EPCAM^-^PDPN^+^LY6C^-^ MHCII^+^) compared to the vehicle, ProAgio and GPH treatments (**Figure 6C**).

**Figure 6:**
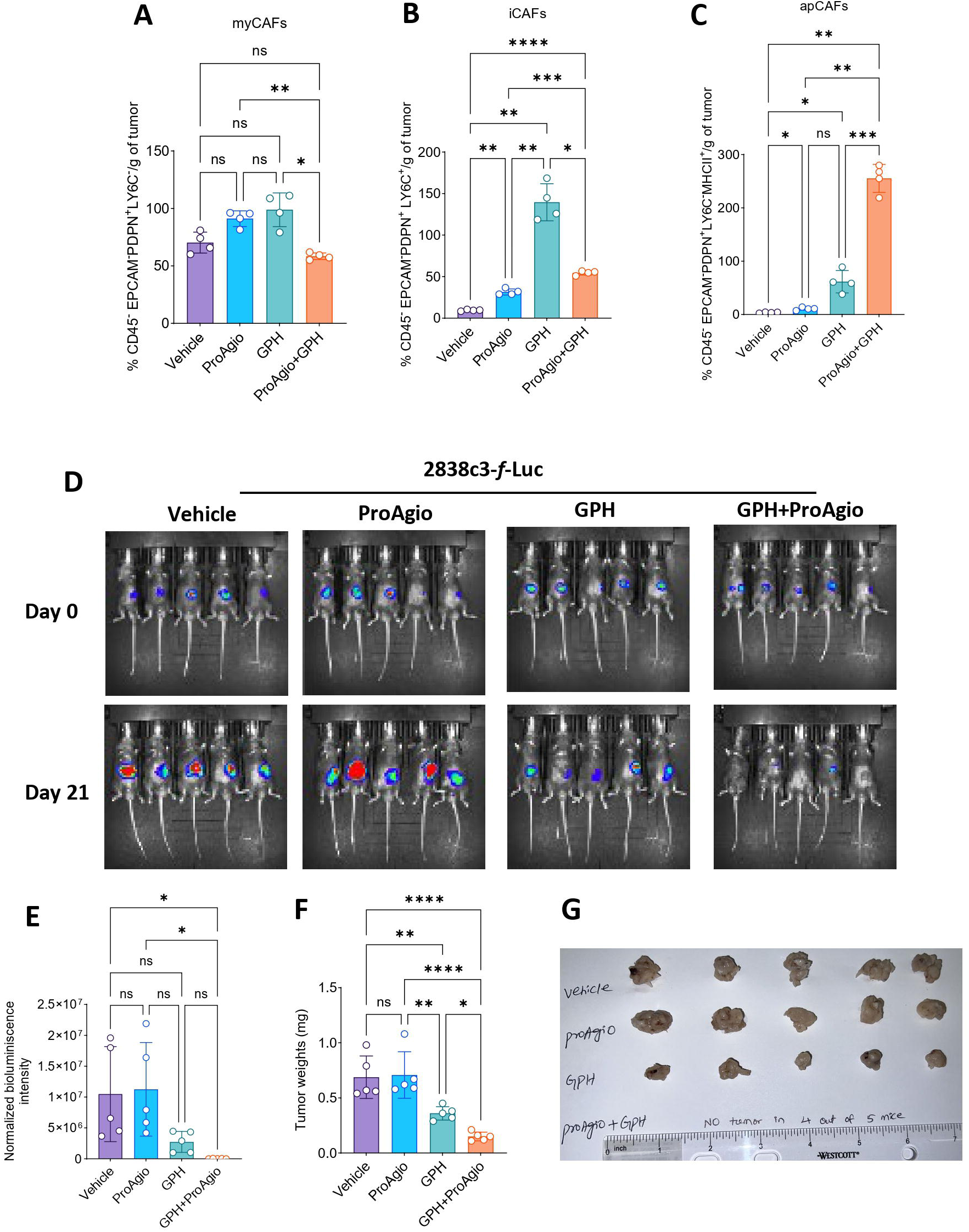
GPH plus ProAgio modulates CAFs and remodels the tumor stromal interactions and inhibits the tumor growth in an immune-rich orthotopic PDAC model. **(A-B).** Reduction in the expression of myCAFs **(A)** and iCAFs **(B)** in GPH plus ProAgio compared to vehicle, ProAgio, and GPH. **(C).** Increased expression of apCAFs in GPH plus ProAgio compared to vehicle, ProAgio, and GPH. **(D).** 2838c3-*f*-luc-labeled cells were orthotopically implanted into C57BL6/J mice (n=20). After 1 week, five mice were orally administered vehicle, ProAgio, GPH, or GPH plus ProAgio. Bioluminescence images at the indicated time points are shown. **(E).** Normalized bioluminescence values from the panel (D). **(F).** Percentage of tumor weights at the indicated conditions for the experiment in (D) **(G).** Tumor images were taken at the end of the experiment for panel (D). Statistical significance was assessed using One-way ANOVA. All quantitative data represent the mean ± SEM. ns, non-significant, ∗*p* < 0.05, ∗∗*p* < 0.01, ∗∗∗*p* < 0.001, and ∗∗∗∗*p* < 0.0001.

### Confirmation of GPH plus ProAgio effects in a second orthotopic mouse model

We then evaluated whether this effect is reproducible in another model. The murine 2838c3-*f*-luc model has more infiltration of T cells at baseline. Murine 2838c3-*f*-luc cells were orthotopically implanted into C57BL6/J mice (n=5 animals/group) and treated with either vehicle, ProAgio, GPH, or GPH plus ProAgio. Interestingly, 4 out of 5 mice treated with the GPH and ProAgio combination had a complete response with no viable tumor (**Figure 6D** and **6E**). In addition to this, we have also identified a significant reduction in the size and weight of the tumors (**Figure 6F** and **6G**). This anti-tumor activity observed in the immune-rich microenvironment suggests that GPH plus ProAgio anti-tumor activity was partly mediated by immune modulation and highlights the potential of this combination in enhancing anti-tumor immunity, possibly involving T-cell activations.

### The combination of GPH plus ProAgio effectively inhibits KRAS^G12D^ PDX growth

Next, we wanted to examine if the effect of GPH plus ProAgio observed in preclinical murine models could be recapitulated in a human patient-derived xenograft (PDX) model of PDAC. This study used a KRAS^G12D^ mutant PDX subcutaneously implanted into an immunocompromised (NSG) mouse. The treatment was initiated after the tumors reached a minimal size of 100 mm³. Mice were randomized and treated with vehicle, ProAgio, GPH, or GPH plus ProAgio. The combination treatment significantly reduced the tumor burden compared to single-agent treatments or vehicle (**Figure 7A-C**). Immunohistochemical analysis showed a significant reduction in the expression of CD31 (**Figure 7D**) and HIF-1α (**Figure 7E**), supporting the previous findings of reduced angiogenesis and hypoxia. In summary, the GPH plus ProAgio combination therapy in the PDX model closely recapitulates the findings observed in the murine PDAC models.

**Figure 7:**
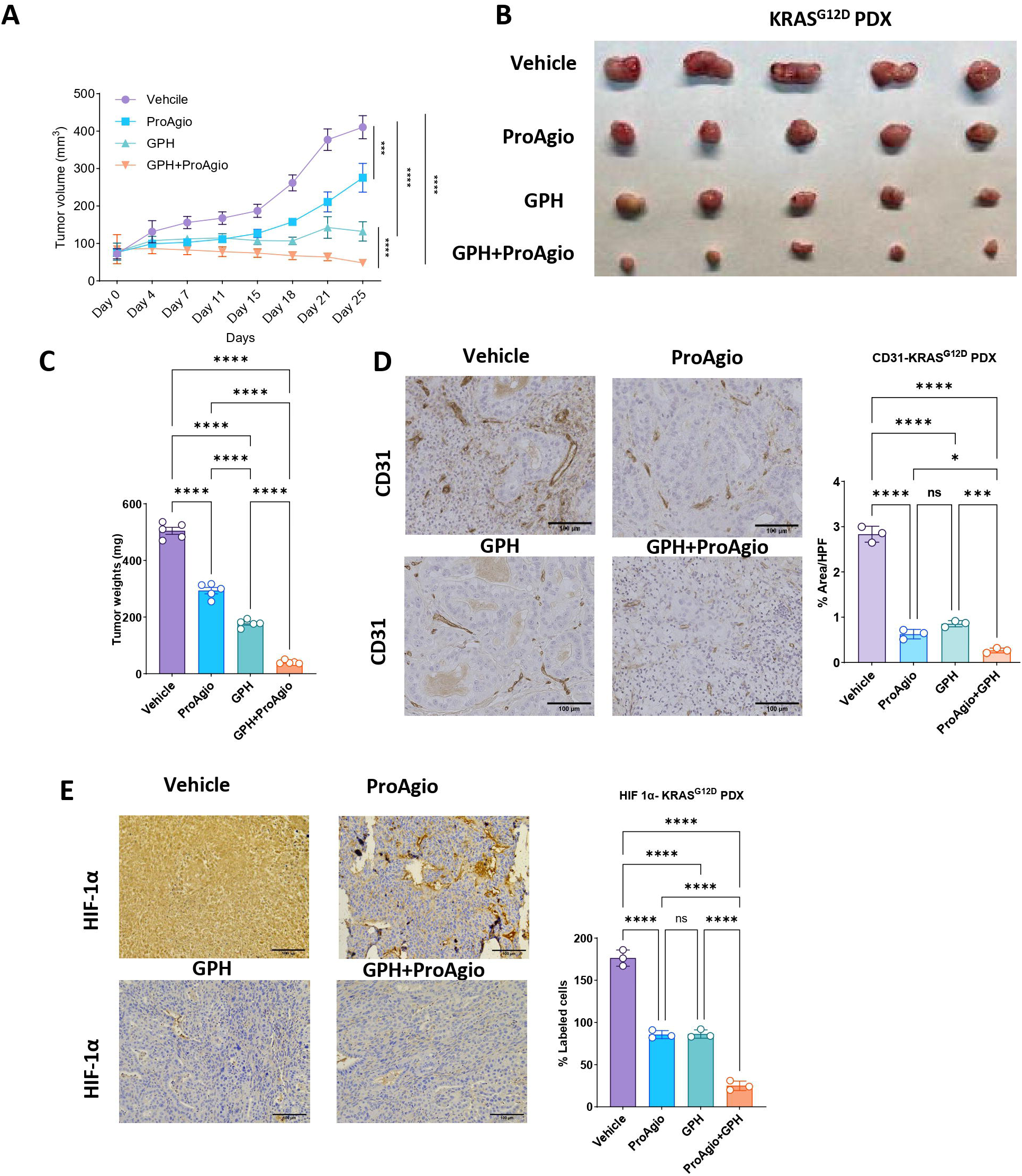
GPH plus ProAgio combination reduces tumor growth and angiogenesis in KRAS^G12D^ PDX models. **(A).** Tumor growth curves of KRAS^G12D^ PDX implanted subcutaneously into NSG mice and treated with vehicle, ProAgio, GPH, or GPH+ProAgio combination (n=5). (**B**). Tumor images showing reduced size in the GPH plus ProAgio combination compared to vehicle, ProAgio, or GPH. (**C**). Percentage of tumor weights at the end of the experiment under the indicated conditions. **(D)**. Representative IHC images showing the reduced expressions of CD31 (left) and quantifications (right) in the GPH plus ProAgio combination compared to vehicle, ProAgio, or GPH. **(E)**. Representative IHC images show the reduced expressions of HIF-1α (left) and quantifications (right) in the GPH plus ProAgio combination compared to vehicle, ProAgio, or GPH. One-way ANOVA and area under the curve (AUC) were used to determine *p*-values. Error bars indicate SD. ns, non-significant, ∗*p* < 0.05, ∗∗∗*p* < 0.001 and ∗∗∗∗*p* < 0.0001. Scale bar 100 µm.

## Discussion

Desmoplasia, tumor hypoxia, and immunosuppressive TME contribute to resistance of PDAC to systemic cytotoxic and immune therapies ^23, 24^. In the current study, we explored the role of integrin α_v_β_3_-expressing endothelial cells and CAFs as central regulators of the TME. The results from this study support the finding that integrin α_v_β_3_ is both a prognostic marker and therapeutic target in PDAC. We evaluated the therapeutic potential of targeting cells expressing integrin α_v_β_3_ by ProAgio with GPH, a regimen known to modulate the TME in PDAC.

Our initial analysis of human tissue array revealed that ITGB3 protein was significantly overexpressed in PDAC tissues compared to normal pancreas, which was further supported by TCGA gene expression data. The increased expression of ITGB3 correlated with markers of angiogenesis, hypoxia, and EMT. Integrin α_v_β_3_plays a key role in several growth-promoting features, such as cell adhesion, migration, and metastasis ^25^. Inhibiting integrin α_v_β_3_ can result in vascular remodeling, impaired invasion and modulation of tumor stiffness. These effects are due to interactions between integrin α_v_β_3_and the ECM ^26–28^. In addition, previous findings also reported that overexpression of ITGB3 resulted in EMT and endothelial to mesenchymal transition (End-MT), which are known hallmarks for PDAC’s aggressive growth, invasiveness, and chemoresistance ^29, 30^. ITGB3 has been shown to promote angiogenesis through TGF-β signaling and contributes to transforming malignant features in cancer cells ^25^. PDAC is known to have leaky blood vessels in the tumor leading to increased tissue interstitial pressures and decreased perfusion with resultant nutrient and oxygen-deprived state ^31^. These conditions lead to the activation of autophagy and promotion of a more invasive and resistant mesenchymal phenotype.

ProAgio has a novel mechanism of action. While other inhibitors block the enzymatic activity of integrin, ProAgio binds to a non-catalytic site, leading to activation of apoptosis. Therefore, ProAgio is a cytotoxin to integrin α_v_β_3_-expressing cells. In our study, we demonstrated that treatment with ProAgio decreased the expression of integrin α_v_β_3_. The combination of ProAgio and GPH had a more pronounced effect on integrin α_v_β_3_-expressing cells than ProAgio alone. The downstream effects of ProAgio include abrogation of the aberrant angiogenesis, which leads to a reduction in vascular leakage, improvement in interstitial pressure, increased perfusion, and reduction in hypoxia within the TME. This study has demonstrated these effects at multiple levels, including tissue-based evaluation of angiogenesis, functional assays to evaluate vascular integrity and leakage and physiological evaluation of blood flow using DCE-MRI. The volume transfer constant (*K^trans^*) measured by DCE-MRI reflects the efflux rate of gadolinium contrast from blood plasma into the extracellular tissue. *K^trans^* depends on vascular permeability, plasma blood flow, and capillary surface area. Based on our models, ProAgio lowers the capillary surface area and vascular permeability. Hence, we postulate that the main driver of the changes in *K^trans^* is the increased flow due to decreased interstitial edema due to lower vascular permeability and improved architecture of vascular channels within the tumor vasculature structure. These effects were also observed in patients treated on the clinical trial at a therapeutic dose of ProAgio, further confirming this mechanism of action. The enhanced tumor blood flow impacts hypoxia in the TME as evidenced by changes in multiple hypoxia markers, such as HIF-1α.

We postulate that this, in turn, inhibits EMT, leading to a less resistant and invasive phenotype. Tumor cells are considered to have undergone EMT, resulting in the loss of epithelial marker expressions in tandem with a gain in mesenchymal marker expressions. Key downregulated epithelial markers include E-cadherin, cytokeratins (such as CK8, CK18, and CK19), and desmoplakin, which were reported in PDAC ^32^. The mesenchymal phenotype has increased N-cadherin, vimentin, S100a4, and α-SMA ^33^ expression. The transition from mesenchymal to epithelial phenotype may reduce the invasive potential of PDAC cells and confer greater sensitivity to chemotherapy and immunotherapy ^34^. In our previous studies, we had shown that GPH can inhibit EMT ^35^. This study shows that the effects on EMT are more significant when we combine GPH with ProAgio. PDAC cells heavily rely on glycolysis as their primary pathway to generate energy to promote growth and survival ^36, 37^. The downregulation of GLUT1, LDHA, and HK1 with the combination therapy may be mediated through inhibition of HIF-1α and its transcriptional target CA9, further supporting the hypothesis that GPH and ProAgio therapy induces metabolic reprogramming of PDAC TME, which impairs cancer cells’ ability to survive the nutrient-deprived microenvironment.

As predicted by our hypothesis, GPH plus ProAgio modulated the TME, shifting these tumors from being immunologically cold to hot. Multi-parameter flow cytometry analysis revealed a significant increase in the CD45^+^ immune cells with a notable increase in γδ T cells, T NK, effector CD4^+^ T-cells, and a shift in the M1/M2 ratio. All these observed effects point to TME reprogramming favoring immune priming and activation. The increased CD4^+^ effector T-cells are crucial in coordinating anti-tumor immunity by producing cytokines and increasing cytotoxic activity ^38^. The observed increase in immune checkpoint markers such as PD-1 and CTLA-4 on T-cells and PD-L1 on macrophages further supports the notion that the combination therapy synergizes with immune checkpoint blockade therapy to induce tumor regression ^39^.

Another target of GPH plus ProAgio is the CAFs, which interact with tumor cells to create a fibrotic environment that promotes chemoresistance and immune evasion ^40–42^. In the current study, we observed a significant reduction in CAFs due to treatment with GPH and ProAgio. Examining the specific CAF phenotypes, we observed that GPH increased iCAFs, and these results are consistent with previous reports showing chemotherapy alone significantly increased the number of CAFs by enhancing the TGFβ signaling pathway ^43^. Adding ProAgio to GPH decreased the iCAF populations, suggesting a potential strategy to counteract therapy-induced tumor-promoting inflammatory effects. Similarly, the addition of ProAgio to GPH reduced myCAFs, leading to the reduction in fibrotic markers such as COL1A1 compared to GPH. We observed an increase in the apCAFs with ProAgio and GPH treatment. Previous reports suggest that apCAFs can drive an increase in CD4^+^ T cells. These fibroblasts promote anti-tumor immunity in a CD4^+^ T cell-dependent manner and induce naïve CD4^+^ T cells in an antigen-specific manner ^44–46^.

GPH has been shown to modulate TME and increase immune activation. Our preclinical and clinical data strongly support the hypothesis that targeting cells that express integrin α_v_β_3_ by ProAgio can potentiate the effects of GPH on the TME in PDAC. ProAgio can modulate tumor angiogenesis to increase blood flow, decrease hypoxia and favor a metabolic switch. These effects contribute to the improved activity of cytotoxic chemotherapy and immune checkpoint blockade. **Figure 8** is a schematic of this combination’s proposed mechanism of action. Gemcitabine and nab-paclitaxel with PH have been evaluated in clinical trials and is safe ^13^. ProAgio is currently in clinical trials with gemcitabine and nab-paclitaxel, and its safety profile appears promising in the early stage. The combination of ProAgio with GPH and nab-paclitaxel is clinically feasible and is supported by the preclinical data presented in this paper. Furthermore, the combination offers a unique multipronged approach to modulate several aspects of PDAC biology that drive resistance and growth. These ProAgio-based combinations are promising and warrant further evaluation in future clinical trials.

**Figure 8:**
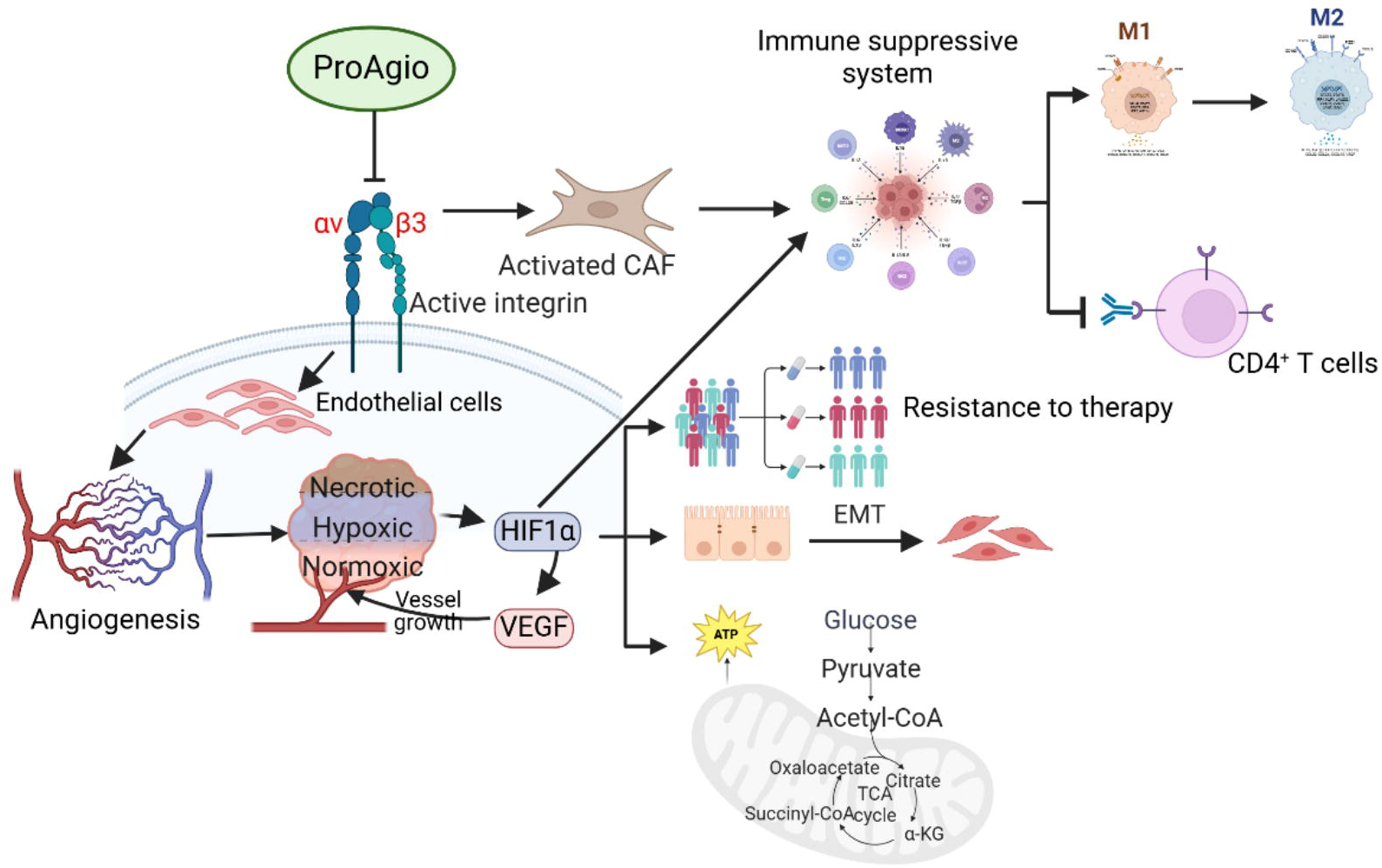
Schematic diagram of the mechanism of action of ProAgio in PDAC.

## Author Contributions

Conceptualization: D.S.R.B., and B.E. Funding acquisition: B.E. Mouse experiments: D.S.R.B., and G.P.N. Immunohistochemistry: D.S.R.B. TCGA data mining: S.S., and D.S.R.B. MRI animal studies: G.P.N., and H. K; Supervision: B.E. Writing original draft: D.S.R.B. review and editing: D.S.R.B, G.P.N., J.B., S.S., M.A., M.M., A.M., H.K., S.B., Z.R.L., and B.E.

## Data and materials availability

All data associated with this study are present in the paper or the Supplementary materials.

## Supplementary figure legends

**Supplementary Figure 1: Effect of GPH plus ProAgio on CD8^+^ T cell subpopulations** Bar graphs represent the percentage changes in CD8^+^ T cells **(A)**, CD8^+^ CTLA-4^+^ (B), CD8^+^ PD-1^+^ **(C)**, and CD8^+^ CTLA-4^+^ PD-1^+^ T cells **(D)**. **(E)**. Expressions of CD8^+^ subset populations, including naïve, pre-effector, effector, and central memory T cells, in vehicle, ProAgio, GPH, and GPH plus ProAgio. One-way ANOVA was used to determine *p*-values. Error bars indicate SD. ns, non-significant, ∗∗p < 0.01, ∗∗∗p < 0.001, and ∗∗∗∗p < 0.0001.

**Supplementary figure 2: (A).** IHC images showed increased CD68 **(A)** and COL1A1 **(B)** expression. IHC quantifications for CD68 **(C)** and COL1A1 **(D)**. One-way ANOVA was used to determine *p*-values. Error bars indicate SD. ns, non-significant, ∗*p* < 0.05, and ∗∗∗*p* < 0.001.

**Supplementary Figure 3:** Gating strategy used for the multiparameter flow cytometry.

## References

1. Sally Á, McGowan R, Finn K, et al. Current and future therapies for pancreatic ductal adenocarcinoma. Cancers 2022;14:2417.

2. Ziani L, Chouaib S, Thiery J. Alteration of the antitumor immune response by cancer-associated fibroblasts. Frontiers in immunology 2018;9:414.

3. Longo V, Brunetti O, Gnoni A, et al. Angiogenesis in pancreatic ductal adenocarcinoma: A controversial issue. Oncotarget 2016;7:58649.

4. Provenzano PP, Cuevas C, Chang AE, et al. Enzymatic targeting of the stroma ablates physical barriers to treatment of pancreatic ductal adenocarcinoma. Cancer cell 2012;21:418–429.

5. Jacobetz MA, Chan DS, Neesse A, et al. Hyaluronan impairs vascular function and drug delivery in a mouse model of pancreatic cancer. Gut 2013;62:112–120.

6. Li J, Wientjes MG, Au JL-S. Pancreatic cancer: pathobiology, treatment options, and drug delivery. The AAPS journal 2010;12:223–232.

7. Guelfi S, Hodivala-Dilke K, Bergers G. Targeting the tumour vasculature: from vessel destruction to promotion. Nature Reviews Cancer 2024;24:655–675.

8. Huang J, Zhang L, Wan D, et al. Extracellular matrix and its therapeutic potential for cancer treatment. Signal Transduction and Targeted Therapy 2021;6:153.

9. Zhang J, Song J, Tang S, et al. Multi-omics analysis reveals the chemoresistance mechanism of proliferating tissue-resident macrophages in PDAC via metabolic adaptation. Cell Reports 2023;42.

10. Turaga RC, Yin L, Yang JJ, et al. Rational design of a protein that binds integrin αvβ3 outside the ligand binding site. Nature Communications 2016;7:11675.

11. Turaga RC, Sharma M, Mishra F, et al. Modulation of cancer-associated fibrotic stroma by an integrin αvβ3 targeting protein for pancreatic cancer treatment. Cellular and Molecular Gastroenterology and Hepatology 2021;11:161–179.

12. Turaga RC, Satyanarayana G, Sharma M, et al. Targeting integrin αvβ3 by a rationally designed protein for chronic liver disease treatment. Communications Biology 2021;4:1087.

13. Nagaraju GP, Saddala MS, Foote JB, et al. Mechanism of enhancing chemotherapy efficacy in pancreatic ductal adenocarcinoma with paricalcitol and hydroxychloroquine. Cell Reports Medicine 2025;6.

14. Yankeelov TE, Luci JJ, Lepage M, et al. Quantitative pharmacokinetic analysis of DCE-MRI data without an arterial input function: a reference region model. Magnetic resonance imaging 2005;23:519–529.

15. Holland MD, Morales A, Simmons S, et al. Disposable point-of-care portable perfusion phantom for quantitative DCE-MRI. Medical physics 2022;49:271–281.

16. Li Z, Tielbeek JA, Caan MW, et al. Expiration-phase template-based motion correction of free-breathing abdominal dynamic contrast enhanced MRI. IEEE Transactions on Biomedical Engineering 2014;62:1215–1225.

17. Vito A, El-Sayes N, Mossman K. Hypoxia-driven immune escape in the tumor microenvironment. Cells 2020;9:992.

18. Yeo CD, Kang N, Choi SY, et al. The role of hypoxia on the acquisition of epithelial-mesenchymal transition and cancer stemness: a possible link to epigenetic regulation. The Korean journal of internal medicine 2017;32:589.

19. Guo M, Liu MYR, Brooks DG. Regulation and impact of tumor-specific CD4+ T cells in cancer and immunotherapy. Trends in Immunology 2024.

20. Gkountidi AO, Garnier L, Dubrot J, et al. MHC class II antigen presentation by lymphatic endothelial cells in tumors promotes intratumoral regulatory T cell–suppressive functions. Cancer immunology research 2021;9:748–764.

21. Kilian M, Sheinin R, Tan CL, et al. MHC class II-restricted antigen presentation is required to prevent dysfunction of cytotoxic T cells by blood-borne myeloids in brain tumors. Cancer Cell 2023;41:235–251. e9.

22. Bawden E, Gebhardt T. The multifaceted roles of CD4+ T cells and MHC class II in cancer surveillance. Current Opinion in Immunology 2023;83:102345.

23. He L, Zhang X, Shi F, et al. Reprograming immunosuppressive microenvironment by eIF4G1 targeting to eradicate pancreatic ductal adenocarcinoma. Cell Reports Medicine 2024;5.

24. Tulamaiti A, Xiao S-Y, Yang Y, et al. ENO1 promotes PDAC progression by inhibiting CD8+ T cell infiltration through upregulating PD-L1 expression via HIF-1α signaling. Translational Oncology 2025;52:102261.

25. Sesé M, Fuentes P, Esteve-Codina A, et al. Hypoxia-mediated translational activation of ITGB3 in breast cancer cells enhances TGF-β signaling and malignant features in vitro and in vivo. Oncotarget 2017;8:114856.

26. Pang X, He X, Qiu Z, et al. Targeting integrin pathways: mechanisms and advances in therapy. Signal Transduction and Targeted Therapy 2023;8:1.

27. Liu A, Liu Y, Li B, et al. Role of miR-223-3p in pulmonary arterial hypertension via targeting ITGB3 in the ECM pathway. Cell Proliferation 2019;52:e12550.

28. Chastney MR, Kaivola J, Leppänen V-M, et al. The role and regulation of integrins in cell migration and invasion. Nature Reviews Molecular Cell Biology 2025;26:147–167.

29. Cheng C, Liu D, Liu Z, et al. Positive feedback regulation of lncRNA TPT1-AS1 and ITGB3 promotes cell growth and metastasis in pancreatic cancer. Cancer Science 2022;113:2986–3001.

30. Zhu C, Kong Z, Wang B, et al. ITGB3/CD61: a hub modulator and target in the tumor microenvironment. American journal of translational research 2019;11:7195.

31. Lugano R, Ramachandran M, Dimberg A. Tumor angiogenesis: causes, consequences, challenges and opportunities. Cellular and Molecular Life Sciences 2020;77:1745–1770.

32. Prieto-García E, Díaz-García CV, García-Ruiz I, et al. Epithelial-to-mesenchymal transition in tumor progression. Medical Oncology 2017;34:1–10.

33. Serrano-Gomez SJ, Maziveyi M, Alahari SK. Regulation of epithelial-mesenchymal transition through epigenetic and post-translational modifications. Molecular cancer 2016;15:1–14.

34. Yang J, Liu Y, Liu S. The role of epithelial-mesenchymal transition and autophagy in pancreatic ductal adenocarcinoma invasion. Cell death & disease 2023;14:506.

35. Nagaraju GP, Saddala MS, Sarvesh S, et al. Paricalcitol plus hydroxychloroquine enhances gemcitabine activity and induces mesenchymal to epithelial transition in pancreatic ductal adenocarcinoma: A single cell RNA-seq analysis. Cancer Letters 2025:217809.

36. Bandi DSR, Sarvesh S, Farran B, et al. Targeting the metabolism and immune system in pancreatic ductal adenocarcinoma: Insights and future directions. Cytokine & Growth Factor Reviews 2023;71:26–39.

37. Reyes-Castellanos G, Masoud R, Carrier A. Mitochondrial metabolism in PDAC: from better knowledge to new targeting strategies. Biomedicines 2020;8:270.

38. Cachot A, Bilous M, Liu Y-C, et al. Tumor-specific cytolytic CD4 T cells mediate immunity against human cancer. Science advances 2021;7:eabe3348.

39. He X, Xu C. Immune checkpoint signaling and cancer immunotherapy. Cell research 2020;30:660–669.

40. Hurtado de Mendoza T, Mose E, Botta G, et al. Tumor-penetrating therapy for β5 integrin-rich pancreas cancer. Nat Commun 12: 1541, 2021.

41. Neuzillet C, Nicolle R, Raffenne J, et al. Periostin-and podoplanin-positive cancer-associated fibroblast subtypes cooperate to shape the inflamed tumor microenvironment in aggressive pancreatic adenocarcinoma. The Journal of Pathology 2022;258:408–425.

42. Yamashita K, Kumamoto Y. CAFs-associated genes (CAFGs) in pancreatic ductal adenocarcinoma (PDAC) and novel therapeutic strategy. International Journal of Molecular Sciences 2024;25:6003.

43. Kim DK, Jeong J, Lee DS, et al. PD-L1-directed PlGF/VEGF blockade synergizes with chemotherapy by targeting CD141+ cancer-associated fibroblasts in pancreatic cancer. Nature communications 2022;13:6292.

44. Song J, Wei R, Liu C, et al. Antigen-presenting cancer associated fibroblasts enhance antitumor immunity and predict immunotherapy response. Nature Communications 2025;16:2175.

45. Huang H, Wang Z, Zhang Y, et al. Mesothelial cell-derived antigen-presenting cancer-associated fibroblasts induce expansion of regulatory T cells in pancreatic cancer. Cancer cell 2022;40:656–673. e7.

46. Dempsey LA. Antigen-presenting CAFs. Nature Immunology 2022;23:645–645.

