## Supplementary figures and images for "ProAgio, a Novel Integrin αvβ3 Targeted Cytotoxin, Suppresses Tumor Growth and Reprograms the PDAC Microenvironment"

### Supplementary Figure 1

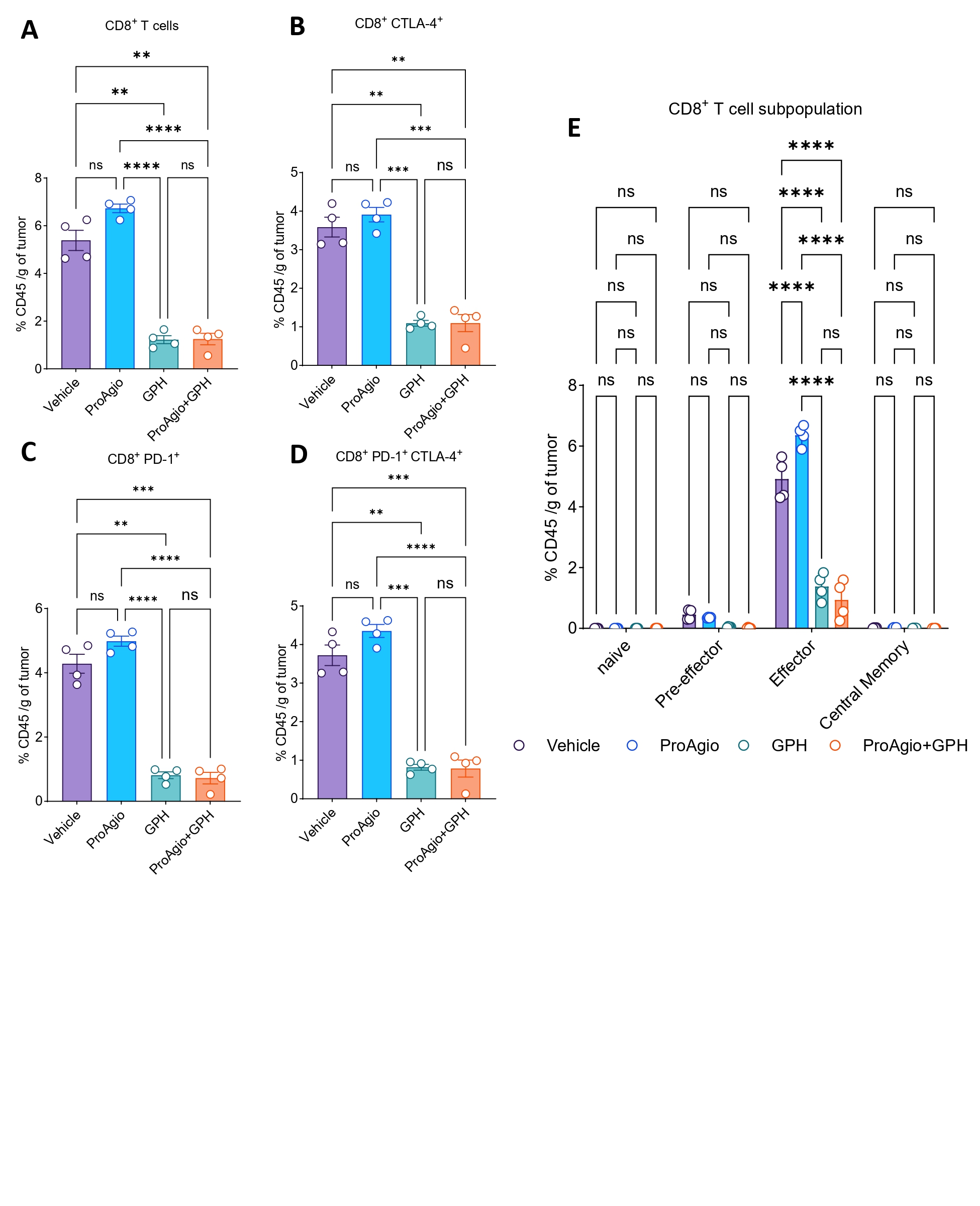

### Supplementary Figure 2

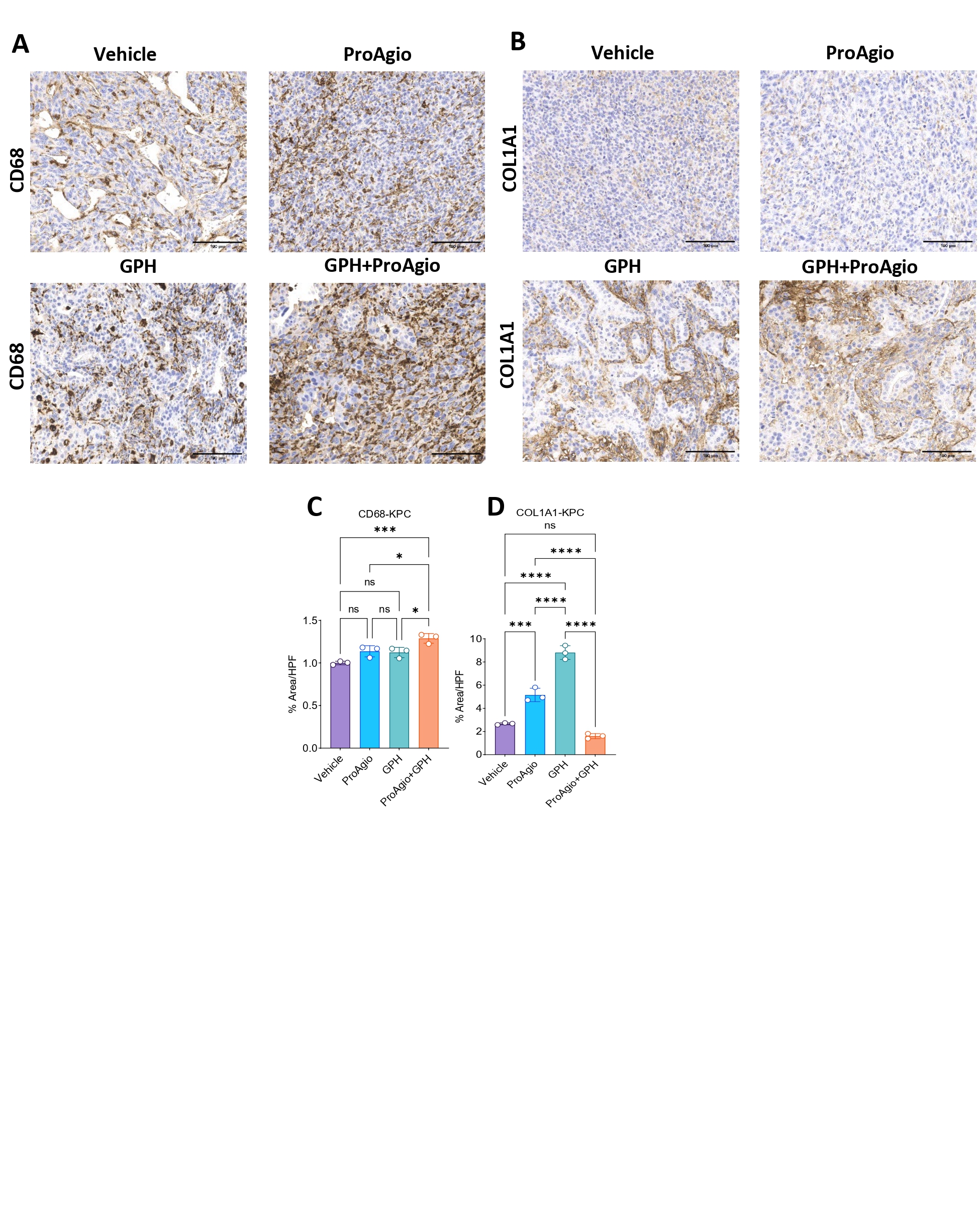

### Supplementary Figure 3

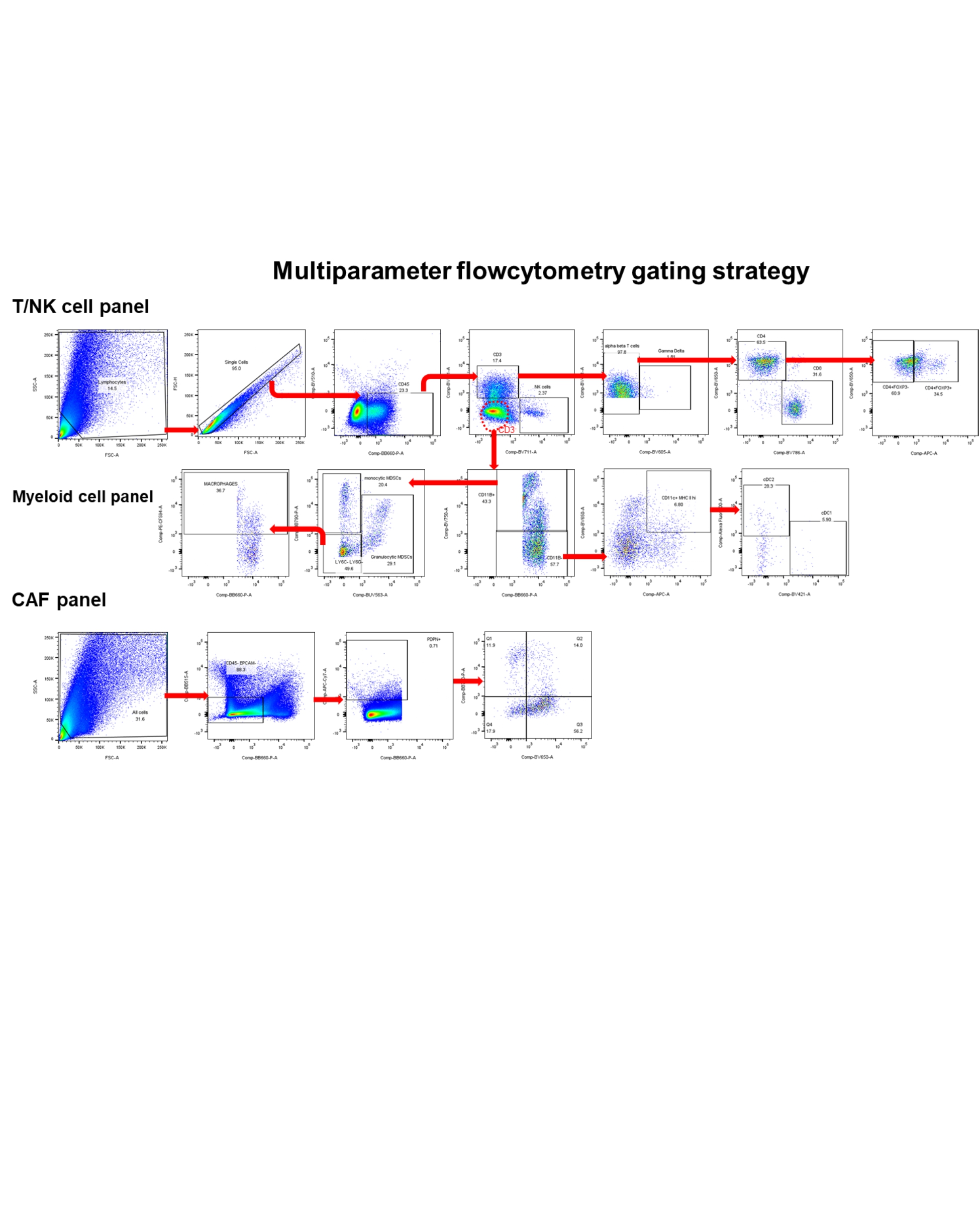
